# SNAP_Switch_: A Molecular Sensor to Quantify the Localization of Proteins, DNA and Nanoparticles in Cells

**DOI:** 10.1101/2019.12.16.877043

**Authors:** Laura I FitzGerald, Luigi Aurelio, Moore Chen, Daniel Yuen, Bim Graham, Angus P R Johnston

## Abstract

Intracellular trafficking governs receptor signalling, pathogenesis, immune responses and the cellular fate of nanomedicines. These processes are typically tracked by confocal microscopy, where colocalization of fluorescent markers implies an interaction or co-compartmentalization. However, this type of analysis is inherently low-throughput, is limited by the resolution of microscopy, and can miss fleeting interactions. To address this, we have developed a localization sensor composed of a quenched and attachable SNAP-tag substrate (SNAP_Switch_). SNAP_Switch_ enables quantitative detection of protein, nucleic acid and nanoparticle trafficking to locations of interest within live cells using flow cytometry. Using this approach, we followed the trafficking of DNA complexes travelling from endosomes into the cytosol and to the nucleus. We also show that antibody targeted to the transferrin (CD71) or hyaluronan (CD44) receptor is initially sorted into different compartments following endocytosis. These results demonstrate SNAP_Switch_ is a high-throughput and broadly applicable tool to quantitatively track the localization of materials in cells.

## Introduction

Determining where material is trafficked following endocytosis is essential for understanding many cellular processes. The intracellular transport of inbound biomolecules is an integral process in cell signalling,^1^ immune responses^2^ and in the manifestation of disease, either through mislocalization of endogenous proteins^3^ or trafficking of infectious agents such as viruses^4^ and bacterial toxins.^5^ Furthermore, the destination of internalized material is of critical importance for the subcellular delivery of therapeutics.^6^ Following uptake, many materials become trapped in endo/lysosomes, preventing access to their site of action.^7,8^ For example, nucleic acids for gene silencing^9^ or delivery^10,11^ require transfer to the cytosol or nucleus respectively. To design carriers that can efficiently deliver molecules to these locations, we need to understand the trafficking of these vehicles and their ultimate subcellular fate.

The intracellular trafficking of materials is commonly investigated through colocalization analysis. The cargo and intracellular compartments of interest are labelled with fluorescent markers such as synthetic organic dyes or fluorescent fusion proteins. The spatial overlap between these fluorescent markers is measured by microscopy and statistical analysis (e.g. Pearson correlation coefficient) is then used to quantify the degree of colocalization.^12^ In certain situations, the interpretation of these results is straightforward, such as when complete colocalization (or alternatively, a complete lack of colocalization) occurs. However, the meaningfulness of intermediate coefficient values between these extremes can be difficult to interpret.^13^ In addition, traditional colocalization analysis is low-throughput, as the number of cells that can be analyzed is limited compared to other techniques such as flow cytometry. Furthermore, the temporal resolution of current techniques means short-lived events, such as trafficking through a particular location may be missed. Spatial resolution is also limited as even super-resolution techniques are limited to >40 nm resolution, which is not sufficient to determine upon which side of a subcellular membrane the material is located.

To identify subcellular structures, fluorescent fusion proteins have traditionally been used to specifically label locations within cells^14^. More recently, genetically encodable tagging systems have attracted attention as an alternative. These techniques, including the biarsenical-tetracysteine tag,^15^ HaloTag,^16^ CLIP-^17^ and SNAP-tag,^18^ were developed to allow fluorescent labelling with small organic dyes which can be superior in terms of brightness and photostability.^19^ Of these, one of the most established is the SNAP-tag, which uses an engineered variant of the 19 kDa O^6^-alkylguanine-DNA alkyltransferase repair enzyme that reacts with benzylguanine, covalently linking a fluorescent label to the protein. Further advances to the SNAP-tag system have been made, such as the advent of quenched substrates, allowing for live cell, wash-free imaging.^20,21^ However, the quenched substrates are engineered to be membrane permeable and are not designed for attachment to DNA, proteins or nanoparticles, which prevents their use in studying the localization of internalized material. To overcome the limitations with conventional co-localization analysis, we have developed a new quenched and attachable SNAP-tag substrate (SNAP_Switch_) for investigating the localization of molecules in live cells. The sensor is attached to a material of interest using click chemistry and remains non-fluorescent until it reaches a SNAP-tagged location of interest within the cell (**Figure 1**). This allows cargo to be detected only when it reaches the location of interest. This is a significant advantage over traditional fluorescent tags, as high-throughput techniques such as flow cytometry can be used to quantify localization, without the need for microscopy and image based colocalization analysis. We have demonstrated the stability and activation of the sensor both in solution and *in vitro*. We then applied this method to probe the differences in trafficking of antibodies bound to cell-surface receptors. We also followed the journey of DNA delivered with a transfection reagent (Lipofectamine) from endosomes, into the cytosol and finally its delivery to the nucleus.

**Figure 1.**
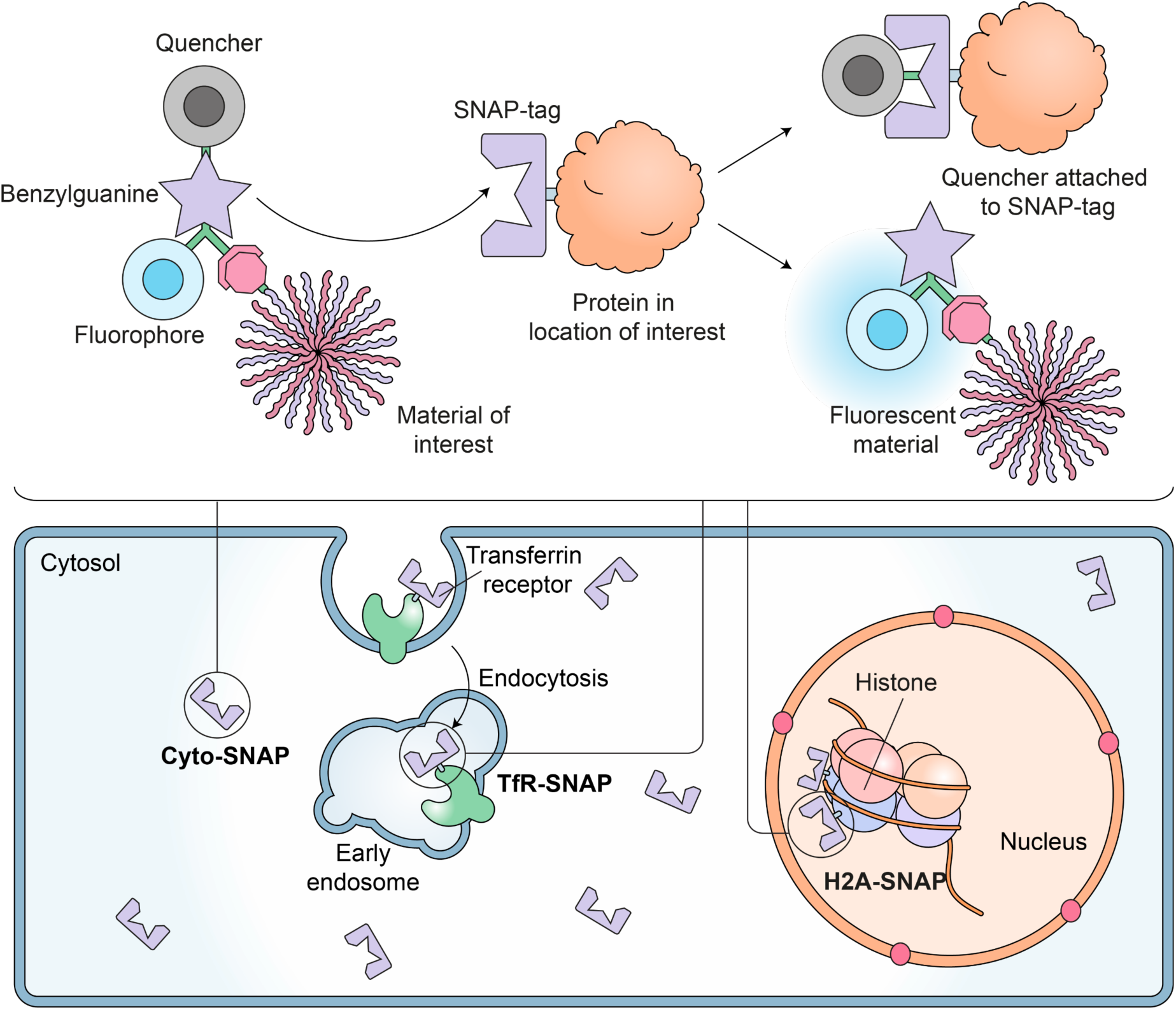
SNAP_Switch_ localization sensor can be used to quantify trafficking of proteins, DNA or nanoparticles to any subcellular compartment. SNAP_Switch_ consists of a quencher (QSY-21) and fluorophore (Cy5) conjugated to either side of a benzyl guanine group. An azide is also included for attachment of the sensor to a material of interest through click chemistry. When the SNAP_Switch_ labelled material reaches a cellular compartment labelled with a SNAP-tag (e.g. early endosome, cytosol or nucleus), the quencher is transferred to the SNAP-tag, breaking the quenching interaction and allowing the sensor to fluoresce.

## Results

### Design of the SNAP_Switch_ Localization Sensor

SNAP_Switch_, is based on benzylguanine, the native substrate for the SNAP-tag. A fluorophore and azido lysine (for subsequent click conjugation to the material of interest) were coupled to the guanine group of benzylguanine, and a quencher was coupled to the benzyl group (**Figure 1, SI Figure 1**). The sensor is engineered so that when SNAP_Switch_ interacts with SNAP-tag, the quencher (QSY-21) is transferred to the SNAP-tag, while the fluorophore (Cy5) becomes fluorescent and remains attached the protein/DNA/nanoparticle. After activation, the fluorescence is permanently switched on. This feature of SNAP_Switch_ offers a significant advantage over image-based colocalization analysis, as signal accumulates over time as more interactions occur. This enables fleeting interactions, such as cargo passing through an endosome into the cytosol to be quantified. Additionally, as the material separates from the SNAP-tag after the interaction, subsequent trafficking of the material can be observed, unlike with assays based on complementation, such as split-GFP, which remains tethered to the protein of interest, potentially blocking further trafficking.

### Sensor Characterization in Solution

Following synthesis, we evaluated the quenching and activation of SNAP_Switch_ by in-gel fluorescence. SNAP_Switch_ exhibited low background fluorescence in its quenched state but in the presence of the SNAP-tag, became intensely fluorescent (**Figure 2a, b**). The fluorescent band of <10 kDa corresponds to the cleaved free dye, and not the molecular weight of the SNAP-tag, as the quencher is transferred to the SNAP-tag and the fluorophore is released. Labelling of free SNAP-tag with the commercial substrate SNAP-Cell SiR 647 resulted in a fluorescent band at ∼20 kDa, while unreacted fluorescent substrate appeared at a similar position to the cleaved SNAP_Switch_ (**Figure 2a**).

**Figure 2.**
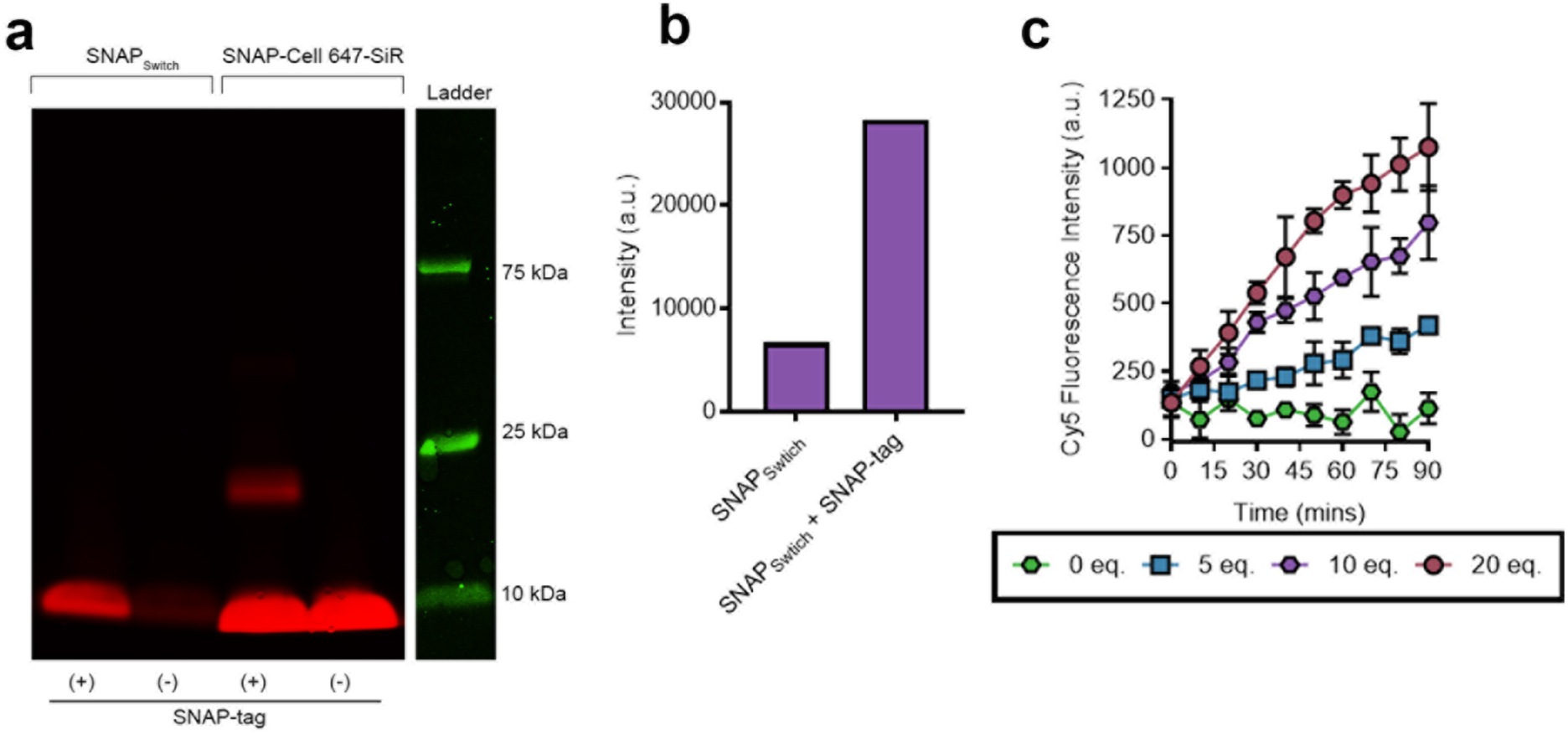
SNAP_Switch_ is specifically activated in the presence of SNAP-tag. (a) In-gel detection of the quenched SNAP-tag substrate of 600 μM SNAP_Switch_ or fluorescent SNAP-Cell 647 SiR with or without 5 μM SNAP-tag protein incubated at 37°C for 30 minutes. (b) Densitometric analysis of SNAP_Switch_ activation with and without SNAP-tag incubation. (c) Fluorescence activation of 0.10 μM SNAP_Switch_ in the presence of SNAP-tag protein at 27°C over a 1.5-hours.

SNAP_Switch_ was stable in the quenched state, showing no increase in fluorescence intensity over a 90-minute incubation period when no SNAP-tag was present. However, the fluorescent signal rapidly activated in the presence of SNAP-tag. Incubation of SNAP_Switch_ with 20 equivalents of SNAP-tag resulted in a ∼10-fold increase in Cy5 fluorescence intensity after 90 minutes (**Figure 2c**). Activation of the sensor was rapid, with a significant difference in SNAP_Switch_ signal compared to the control observed within 10 minutes.

### *In Vitro* Activation of SNAP_Switch_

Having established that SNAP_Switch_ was quenched efficiently and was responsive to SNAP-tag in solution, we moved to demonstrate its response *in vitro*. Activation of SNAP_Switch_ by SNAP–tag expressed in cells was tested by fusing SNAP-tag to the human transferrin receptor (hTfR-SNAP). hTfR-SNAP was stably introduced into NIH/3T3 (3T3) cells using lentiviral transduction and was expected to localize with endogenous mouse TfR. SNAP_Switch_ was then conjugated to either anti-mouse TfR (anti-mTfR) or anti-human TfR (anti-hTfR) antibodies. In contrast to holo-transferrin which rapidly recycles back to the surface, these antibodies are expected to be trafficked to the lysosome over time.^22^

To determine if SNAP_Switch_ could be activated non-specifically, we also conjugated the sensor to an anti-CD44 antibody. CD44 is a membrane glycoprotein expressed by 3T3^23^ cells that is involved in cell adhesion, signaling and is predominately located on the plasma membrane.^24^ An additional fluorophore, BODIPY FL (BDP-FL) was also attached to the antibodies to detect the proteins while the sensor was switched off. This also allowed us to account for slight variations in the amount of association by calculating the ratio of the Cy5 signal generated by the SNAP_Switch_ to the amount of antibody present determined by the BDP-FL signal.

SNAP_Switch_ was specifically activated by proteins that colocalized with hTfR-SNAP. As expected, anti-CD44 and anti-mTfR associated with both wild type and hTfR-SNAP expressing 3T3 cells (**Figure 3a**). Anti-hTfR bound only to cells expressing the hTfR-SNAP fusion protein, confirming the specificity of anti-hTfR for human TfR. SNAP_Switch_ was not activated in cells lacking the SNAP-tag, as no Cy5 signal was observed for any antibody in wild type 3T3 cells after 1 hour (**Figure 3b**), and no increase in SNAP_Switch_ signal was observed over 4 hours (**Figure 3c**). This demonstrates the *in vitro* stability and specificity of the sensor. In cells that express hTfR-SNAP, SNAP_Switch_ was activated when conjugated to anti-mTfR (∼400 a.u.) and anti-hTfR (∼900 a.u.) (**Figure 3b**). Activation occurred rapidly, with the majority of signal generated in the first hour, and only a slight increase thereafter (**Figure 3d**). The activation of SNAP_Switch_ attached to anti-mTfR by SNAP-tag fused to hTfR shows that the two isotypes of the receptor colocalize, and that their proximity is such that the SNAP-tag has access to the sensor. This was confirmed by colocalization of fluorescently labelled anti-hTfR and anti-mTfR (**SI Figure 2**). Anti-mTfR and anti-hTfR also colocalized with hTfR-SNAP labelled with a cell permeable fluorescent SNAP substrate (**Figure 3e, f**).

**Figure 3.**
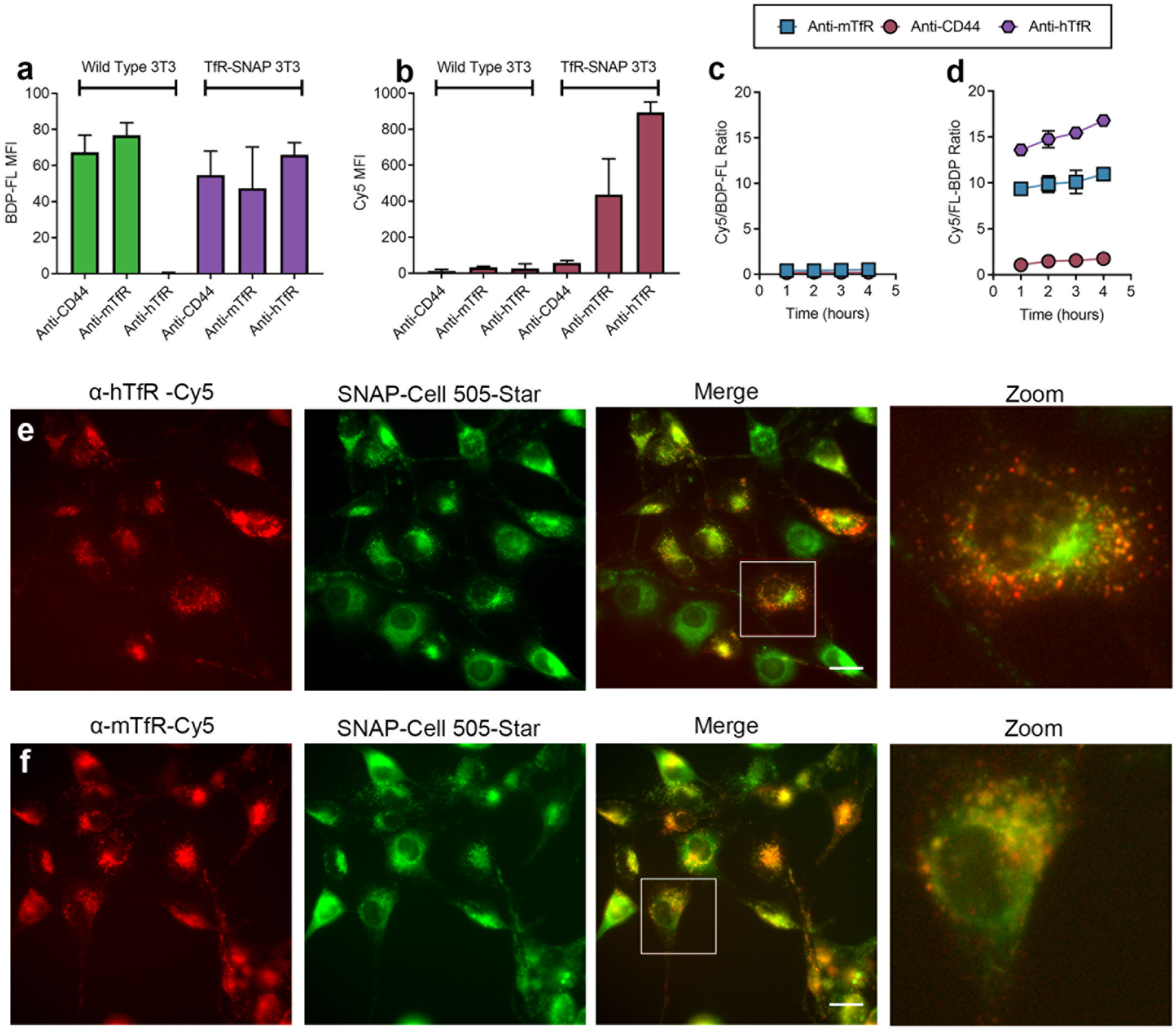
SNAP_Switch_ is activated *in vitro* only when it comes into direct contact with a SNAP-tagged protein. SNAP_Switch_ conjugated to anti-transferrin antibodies is activated by SNAP-tag fused to the transferrin receptor (TfR-SNAP). (a) Association of BDP-FL labelled anti-CD44, anti-mTfR (anti-mouse TfR) and anti-hTfR (anti-human TfR) at 1 hour in 3T3 or 3T3 cells stably expressing SNAP-tag fused to the human TfR (TfR-SNAP), measured by flow cytometry. (b) Activation of SNAP_Switch_ on antibodies in wild type 3T3 or TfR-SNAP cells after 1 hour. SNAP_Switch_ activation on antibodies over time in (c) wild type or (d) TfR-SNAP cells, measured by the ratio of Cy5 to BDP-FL mean fluorescence intensity via flow cytometry at each time point. Fluorescence microscopy images of Cy5 labelled (e) Cy5 labelled (red) anti-hTfR and (f) anti-mTfR in TfR-SNAP cells with the SNAP-tag stained with SNAP-Cell 505-Star (green). (g) Densitometric analysis of SNAP_Switch_ activation, in-gel conjugated to anti-mTfR and treated with 0 – 50 equivalents of SNAP-tag for 1 hour at 37°C, relative to the degree of labelling. anti-mTfR dual-labelled with AF488 and SNAP_Switch_ incubated for 1 hour at 37°C with or without 30 equivalents of SNAP-tag before incubation in either 3T3 or TfR-SNAP cells and analyzed by flow cytometry. (h) Association measured by the AF488 signal. (i) SNAP_Switch_ activation determined by the Cy5/AF488 ratio and (j) the percentage of SNAP_Switch_ activated on anti-mTfR in TfR-SNAP cells. The mean fluorescence intensity, ratio or percentage is plotted with error bars representing the standard deviation of two experiments in triplicate (n = 6). n.s. = non-significant, one-way ANOVA. Scale bar = 20 μm.

In contrast to the anti-TfR antibodies, minimal activation of SNAP_Switch_ attached to anti-CD44 was observed (∼30 a.u.) over 1 hour (**Figure 3b**) even through there was significant association with cells (**Figure 3a**). This suggests anti-CD44 was not trafficked to compartments containing the hTfR-SNAP fusion over this time period. This was corroborated by fluorescence microscopy which demonstrated the majority of anti-CD44 remained bound to the cell surface and did not colocalize with anti-hTfR or anti-mTfR after 1 hour (**SI Figure 3**). However, a small amount of activation did occur over 4 hours (**Figure 3d**), indicating that small amounts of anti-CD44 may eventually reach endocytic vesicles bearing TfR-SNAP. Following a 4-hour incubation of both anti-CD44 and anti-hTfR, punctate structures with both antibodies were visible (**SI Figure 4**).

**Figure 4.**
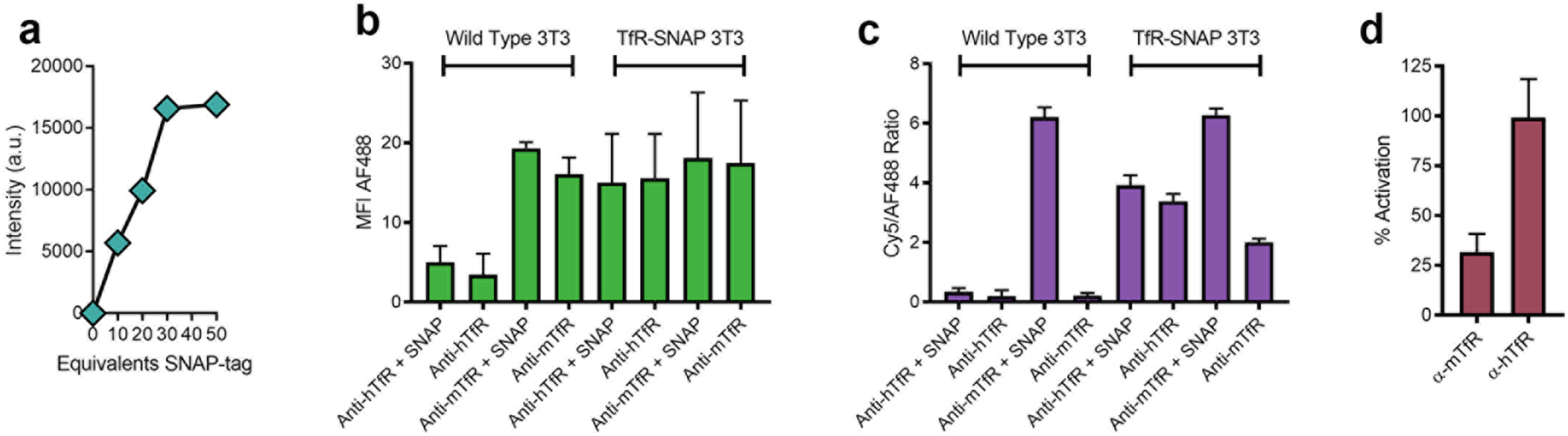
Quantification of SNAP_Switch_ interactions. (a) Densitometric analysis of SNAP_Switch_ activation, in-gel conjugated to anti-mTfR and treated with 0 – 50 equivalents of SNAP-tag for 1 hour at 37°C, relative to the degree of labelling. anti-mTfR dual-labelled with AF488 and SNAP_Switch_ incubated for 1 hour at 37°C with or without 30 equivalents of SNAP-tag before incubation in either 3T3 or TfR-SNAP cells and analyzed by flow cytometry. (b) Association measured by the AF488 signal. (c) SNAP_Switch_ activation determined by the Cy5/AF488 ratio and (d) the percentage of SNAP_Switch_ activated on anti-mTfR in TfR-SNAP cells. The mean fluorescence intensity, ratio or percentage is plotted with error bars representing the standard deviation of two experiments in triplicate (n = 6).

### Quantification of SNAP_Switch_ interactions

The total amount of antibody that interacts with hTfR-SNAP can be quantified by ratioing the SNAP_Switch_ activation in live cells with signal from an antibody that has the SNAP_Switch_ fully pre-activated before incubation with cells. Treating SNAP_Switch_ antibody conjugates (also labelled with AF488 as an unquenched control) with a 30-molar excess of SNAP-tag for 1 hour fully activated the SNAP_Switch_ (**SI Figure 5, Figure 4a**). SNAP-tag pre-treatment did not affect association of the antibodies with either wild type 3T3 or hTfR-SNAP cells (**Figure 4b**). Complete pre-activation of the sensor was confirmed as the SNAP_Switch_:AF488 ratio for anti-mTfR treated with SNAP-tag was the same in both wild type (∼6.2) and hTfR SNAP-tag cells (∼6.3) (**Figure 4c**). To measure the *in vitro* activation of SNAP_Switch_, and thus quantify localization, the ratio of the signal from SNAP_Switch_ activated by hTfR-SNAP to the signal from pre-activated SNAP_Switch_ was calculated. This established that ∼100% of anti-hTfR was activated upon binding to hTfR-SNAP (**Figure 4d**), demonstrating that SNAP_Switch_ is efficiently switched on when it comes into direct contact with the SNAP-tag. In comparison, anti-mTfR was ∼32% activated by hTfR-SNAP, indicating that only a portion of anti-mTfR antibody came into close proximity to the hTfR.

### Trafficking, Endosomal Escape and Nuclear Delivery of Oligonucleotide

We extended the ability of SNAP_Switch_ to track cargo through the endocytic pathway to include trafficking to other locations of interest within the cell. Delivery of nucleic acids to the cytosol and nucleus is critical for gene therapy applications, but the efficiency of this transport is difficult to determine. Lipofectamine 3000, a cationic lipid formulation that condenses DNA into nano-sized complexes,^25^ is widely used as a transfection reagent that induces endosomal escape and facilitates delivery to the nucleus.

To probe the delivery of DNA to the nucleus using Lipofectamine, we fused the SNAP-tag to the nuclear protein histone H2A (H2A-SNAP). A fluorescent protein, mTurquoise was also fused to the H2A-SNAP construct (H2A-mTurq-SNAP) to enable easy identification of SNAP-tagged cells. Fluorescence microscopy confirmed localization of H2A-mTurq-SNAP in the nucleus (**SI Figure 6**).

Cells were transiently transfected with H2A-mTurq-SNAP and delivery of the DNA cargo to the nucleus was evaluated using flow cytometry by measuring the SNAPSwitch signal in live cells positive for mTurquoise. Two populations were identified in the mTurquoise channel (**SI Figure 7**), one with high mTurquoise fluorescence (MFI ∼ 100 a.u.) and one with no expression (MFI ∼ 3 a.u.). The expression of SNAP-tag in the mTurquoise positive population was confirmed by the high SNAP-Cell SiR 647 signal (> 8000 a.u.) within these cells (**Figure 5a**). Dual-labelled Lipofectamine/DNA complexes were then formulated using a 20-mer oligonucleotide sequence labelled with AF488 and SNAP_Switch_. No difference in association of the complexes with wild-type or H2A-mTurq-SNAP HEK cells was observed (**Figure 5b**), allowing for direct analysis of the SNAP_Switch_ signal to determine nuclear access of the delivered DNA (**Figure 5c**). Background signal from the SNAP negative cells showed a small increase in signal over 16 hours, which was subtracted from the Cy5 signal from the mTurquoise positive cells. The SNAP_Switch_ signal in H2A-mTurq-SNAP cells was not significantly higher than in wild type cells over the first 2 hours, suggesting no exposure of the delivered DNA to nuclear components during this period. From 2 to 4 hours, the SNAP_Switch_ signal increased slightly followed by a steady increase in intensity up to 16 hours. This shows that, as expected, there is a lag time between the Lipofectamine/DNA complex being added to the cells and meeting nuclear contents. It is possible that the DNA cargo comes in contact with the SNAP-tagged H2A during mitosis (when the nuclear membrane brakes down) and that direct delivery of DNA through the nuclear membrane does not occur.26 Regardless of the mechanism, SNAP_Switch_ is able to determine if and when the delivered DNA can access the nuclear components of the cell.

**Figure 5.**
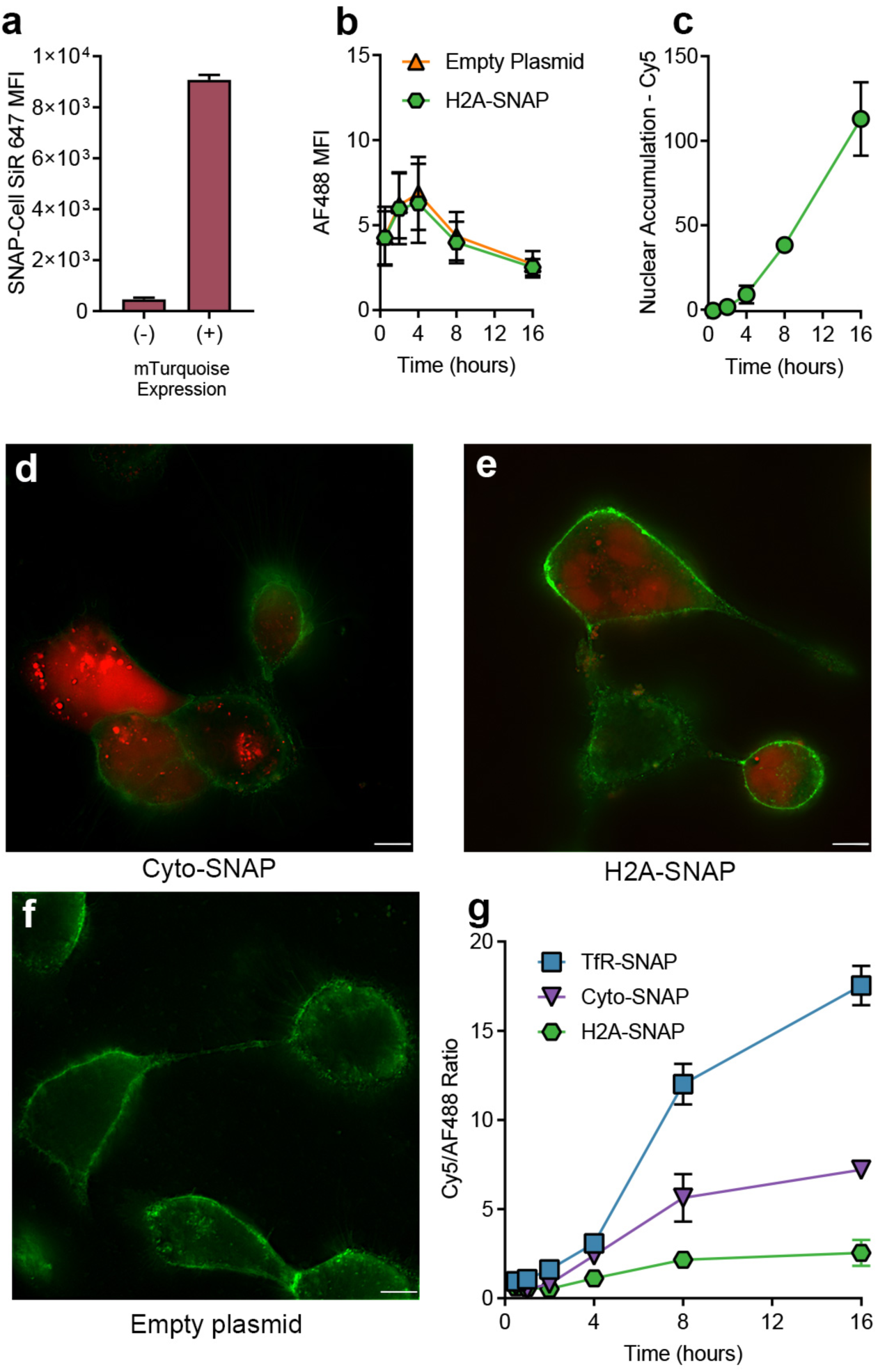
SNAP_Switch_ can follow trafficking, endosomal escape and nuclear transport of oligonucleotides over time. HEK293 cells were transfected to transiently express H2A-mTurquoise-SNAP-tag and analyzed by flow cytometry. (**a**) SNAP-Cell SiR 647 signal in cells gated positive or negative for mTurquoise expression. (**b**) Association of Lipofectamine 3000 complexes with cells transfected with H2A-mTurquoise-SNAP or an empty plasmid. (**c**) SNAP_Switch_ signal in cells positive for mTurquoise expression with the signal from negative cells subtracted as background. Deconvolved fluorescence microscopy images of oligonucleotide labeled with SNAP_Switch_ delivered into cells transfected with (**d**) Cyto-SNAP, (**e**) H2A-SNAP and (**f**) an empty plasmid, scale bar = 10 μm. (**g**) HEK cells stably expressing TfR-SNAP, Cyto-SNAP or H2A-SNAP and transfected with Lipofectamine 3000 complexed oligonucleotides labelled with both AF488 and SNAP_Switch_, the ratio of the SNAP_Switch_ to AF488 signal of oligonucleotide Lipofectamine 3000 complexes with cells over time by flow cytometry. The mean fluorescence intensity or ratio is plotted with error bars representing the standard deviation, in triplicate (n = 3).

To further probe the trafficking of the Lipofectamine/DNA complexes, we also engineered cells to express a cytosolic SNAP-tag (Cyto-SNAP) to measure endosomal escape. Fluorescence microscopy confirmed a diffuse signal throughout the cells for the Cyto-SNAP construct (**SI Figure 8**). To confirm Cyto-SNAP localized in the cytosol and not in lysosomes, we looked for activation of SNAP_Switch_ conjugated to anti-mTfR. We22, and others, have previously shown anti-TfR antibodies localize to endo/lysosomes.27 SNAP_Switch_ labelled anti-mTfR showed no activation in Cyto-SNAP transfected cells after 4 hours but induced a robust response in hTfR-SNAP cells (**SI Figure 9**). This demonstrates that Cyto-SNAP does not localize to the lysosomes. We then used the SNAP_Switch_ sensor to visualize both cytosolic delivery and nuclear exposure by fluorescence microscopy. Cyto-SNAP or H2A-SNAP cells were incubated with SNAP_Switch_ Lipofectamine/DNA complexes for 16 hours.

The response from the sensor was localized to different regions of the cell in those expressing Cyto-SNAP or H2A-SNAP. In Cyto-SNAP cells, SNAP_Switch_ signal was observed as a combination of bright punctate structures as well as dispersed fluorescence throughout the cell (**Figure 5d**). As Cyto-SNAP does not localize to the lysosomes, the punctate structures suggest that a proportion of the DNA may still be complexed with Lipofectamine 3000 in the cytosol or is associated with the membrane remnants of ruptured endo/lysosomes. The fluorescent signal from H2A-SNAP cells was confined to the nucleus (**Figure 5e**). The reduced amount of punctate structures suggests the DNA has dissociated from Lipofectamine either before or after contact with H2A. No signal was observed in wild type cells incubated with SNAP_Switch_ Lipofectamine/DNA complexes and transfected with an empty plasmid (**Figure 5f**).

To demonstrate the high-throughput potential of SNAP_Switch_ using flow cytometry, we further probed the trafficking of the Lipofectamine/DNA complexes over time. Cell lines stably expressing TfR-SNAP, Cyto-SNAP and H2A-SNAP were generated using lentiviral transduction of HEK293 cells. The behavior of the dual-labelled oligonucleotide Lipofectamine 3000 complexes was then followed over 16 hours. The association of the complexes across the cell lines was similar (**SI Figure 10a, SI Figure 11a**). Again, to account for the slight variations in cell association, the ratio of SNAP_Switch_ to AF488 signal at each time point was calculated. The SNAP_Switch_ signal for TfR-SNAP cells showed a small increase in intensity over the first 2 hours, whereas the Cyto-SNAP and H2A-SNAP showed minimal activation (**Figure 4g, SI Figure 10b, SI Figure 11b**). From 4 to 16 hours the TfR-SNAP cells showed significant activation, indicating that lipoplexes are trafficked into TfR positive vesicles. As expected, the signal from the Cyto-SNAP cells was significantly lower than the TfR-SNAP cells after 16 hours, indicating that lipoplexes get trapped in TfR positive vesicles and the endosomal translocation is relatively inefficient. Finally, the H2A-SNAP cells showed the lowest activation, indicating that efficiency of transport from the cytosol to nuclear constituents is also inefficient.

## Discussion

The uptake and intracellular trafficking of material is important for regular cellular function, the mechanisms behind disease pathogenesis and drug delivery. However, current techniques to determine localization are limited by low-throughput, are open to subjective interpretation, and can miss momentary interactions. Through synthesis of a quenched and attachable SNAP-tag substrate, we have developed a sensor that enables high-throughput and quantitative tracking of biomacromolecules in live cells. SNAPSwitch permanently switches on when labelled material encounters SNAP-tagged proteins in specific subcellular locations. This is a significant improvement over other systems such as split-GFP^28^ where both parts must remain bound for signal to occur, which blocks subsequent protein trafficking. It is also a significant improvement over image-based colocalization assays, which struggle to detect transitionary processes such as those that occur during endosomal trafficking and endosomal escape. The use of flow cytometry means that robust statistical analysis of over >10,000 cells can be performed, rather than performing image-based colocalization on <100 cells.

Using SNAP_Switch_, we have demonstrated the ability to quantify trafficking of cell surface receptors, such as TfR (CD71) and CD44. We show that while antibody against CD44 mostly localizes to the cell membrane, small amounts do meet TfR positive compartments over time. The lack of initial SNAP_Switch_ activation attached to anti-CD44 highlights two points: first, even though both hTfR-SNAP and anti-CD44 are present on the surface of the cell, the SNAP_Switch_ is not activated. This indicates that hTfR-SNAP and bound anti-CD44 do not come close enough on the surface of the cell for an interaction to occur. Second, the trafficking pathway followed by anti-CD44 is different to that of the two anti-TfR antibodies. CD44 has been reported to internalize through clathrin-independent carrier (CLIC) endocytosis, rather than the clathrin dependent pathway followed by the transferrin receptor.^29^ The partial activation after further incubation suggests that a proportion of anti-CD44 does reach eventually reach the same compartments as the TfR-SNAP fusion. In addition, we also tracked the uptake of DNA complexed with Lipofectamine into endosomes, followed by its translocation into the cytosol and subsequent trafficking to nuclear components in live cells. We show the amount present in each of these locations decreases along this journey which supports the current view that endosomal escape is a major bottleneck in delivery of nucleic acids.^30^

SNAP_Switch_ provides a new method to follow the journey of material in live cells to subcellular locations following endocytosis. The low background fluorescence of the sensor when in its off state and the large increase when the SNAP-tag is encountered provides the basis for a system to definitively identify when internalized cargo has reached a specific location in the cell. As the sensor is only activated if it comes into close proximity with the SNAP-tag, co-trafficking of proteins and receptors can be investigated. Additionally, the ability to detect delivery to subcellular components will provide a useful tool in the biological sciences to study the intracellular trafficking, endosomal escape and the subcellular delivery of nanoparticles, proteins and nucleic acids.

## Supporting information

Supporting Information figures are available here.

## Supporting Information

Supporting Information figures are available here.

## Acknowledgements

The authors thank Dr. Jason Dang (Monash University, Melbourne) for his assistance with HPLC, LCMS and HRMS.

## Author Contributions

L.I.F., A.P.R.J. and B.G. designed the study. L.I.F, L.A and B.G. performed the chemical synthesis. M.C. and D.Y. performed the cloning. L.I.F. and A.P.R.J. analyzed the data and wrote the manuscript.

## Methods

### Materials

Peptide synthesis reagents including 2-chlorotrityl chloride resin, N(α)-Fmoc-N(ε)-azide-L-lysine and (Benzotriazol-1-yloxy)tripyrrolidinophosphonium hexafluorophosphate (PyBOP) were purchased from ChemImpex. Fmoc-N-amido-dPEG_4_-acid was purchased from Quanta Biosciences. Sulfonated Cyanine 5 succinimidyl ester (Cy5-NHS) and BODIPY-FL succinimidyl ester (BDP-FL-NHS) were purchased from Lumiprobe. Solvents and other organic synthesis reagents including acetonitrile (MeCN), ethylenediamine, dichloromethane (DCM), dimethylformamide (DMF), methanol (MeOH), N,N-diisopropylethylamine (DIPEA), anhydrous dimethyl sulfoxide (DMSO), hexafluoroisopropanol (HFIP), triisopropylsilane (TIPS), Dithiothreitol (DTT), piperidine and phosphate buffered saline tablets (PBS) were purchased from Sigma-Aldrich. Sulfonated QSY-21 carboxylic acid (sQSY-21-COOH)^31^ and Fmoc-benzylguanine carboxylic acid (Fmoc-BG-COOH)^32^ were synthesized and purified in house using published procedures.

Fluorescent SNAP-tag substrates and the SNAP-tag plasmid, pSNAPf were purchased from New England Biolabs. Additional labels and reagents including Alexa Fluor 488 succinimidyl Ester (AF488-NHS), Alexa Fluor 488 Azide, Wheat Germ Agglutinin Alexa Fluor 488 conjugate, Click-IT Succinimidyl Ester DIBO Alkyne (DIBO-NHS), Zeba Spin Columns 7K Molecular Weight Cut-Off, Lipofectamine 3000 Transfection Reagent, Hoechst 33342 Trihydrochloride Trihydrate and general tissue culture supplies were obtained from Thermo Fisher Scientific.

The monoclonal IgG1 anti-mouse CD44 (5035-41.1D) was purchased from Novus Biologicals. Purified mouse monoclonal IgG1 anti-human transferrin receptor antibody (clone OKT9)^33^ and anti-mouse transferrin receptor IgG2a monoclonal antibody (clone TIB-219)^34^ were purchased from Antibody Services at the Walter and Eliza Hall Institute Biotechnology Centre.

### Instrumentation

Liquid Chromatography – Mass Spectrometry (LCMS) was performed on an Agilent 6100 Series Single Quad LCMS with a photodiode array detector (214/254 nm) coupled to an Agilent 1200 Series HPLC with a G1311A quaternary pump, G1329A thermostated auto sampler and 1200 Series G1314B variable wavelength detector with a scan range between 100–1000 m/z and a 5-minute acquisition time.

High-performance Liquid Chromatography (HPLC) was performed on an Agilent 1260 series modular HPLC fitted with a G1312B binary pump, G1316A compartment equipped with an Agilent Eclipse Plus C18 3.5 µm, 4.6 x 100 mm column and a G1312B diode array detector using an elution protocol of 0 – 10 min, gradient from 5% MeCN/0.1% TFA/95% H_2_O/0.1% TFA to 100% MeCN/0.1% TFA with a flow rate of 1 mL min^-1^. Preparative high-performance liquid chromatography (HPLC) used a Grace Alltima C8 5μ particle size, 22 x 250 mm column.

High Resolution – Mass Spectrometry (HRMS) was performed on a Waters LCT TOF LCMS Mass Spectrometer coupled to a 2795 Alliance Separations module. All data was acquired, and mass corrected via a dual-spray Leucine Enkephaline reference sample. Mass spectra were created by averaging the scans across each peak and background subtracted of the TIC. Acquisition and analysis were performed using the Masslynx software version 4.1.

Gel fluorescence images were obtained on an Amersham Typhoon 5 Biomolecular Imager (GE Healthcare Life Sciences) and analyzed in ImageJ. Fluorescence intensity measurements in solution were performed with a PerkinElmer EnSpire Multilabel plate reader operating at 27°C. All absorbance measurements were obtained with a NanoDrop ND-1000.

### Cell Culture

NIH/3T3 (ATCC: CRL-1658) and HEK293A (HEK, ThermoFisher Scientific R70507) were maintained in Dulbecco’s Modified Eagle Medium (DMEM), high glucose (GlutaMAX) with phenol red and 20% (NIH/3T3) or 10% (HEK293) fetal bovine serum (FBS) and 1% penicillin/streptomycin at 37oC with 5% CO_2_.

### Flow Cytometry

10,000 events per sample were analyzed with a Stratedigm S1000EXI flow cytometer (Stratedigm, California, USA) using 405 and 552 nm excitation with emission collected between 415 and 475 nm (for mTurquoise), 488 nm excitation with emission collected between 515 – 545 nm (for Alexa Fluor 488/BDP-FL) and 642 nm excitation with emission collected between and 661 – 690 nm (for Cy5/SNAP-Cell SiR 647). FCS3.0 files were exported using CellCapTure Analysis Software (version 4.0 RC4, Stratedigm, California, USA) and imported into FlowJo (version 8, Tree Star, Oregon, USA).

For analysis, untreated cells were gated on the forward versus side scatter log area plot and the gate was then applied to all samples (**SI Figure 12**). The geometric mean fluorescence intensity of untreated cells was taken as the background. The average of this was then subtracted from each sample. These values were then plotted directly or the ratio of Cy5 to BDP-FL or AF488 in each well was calculated. The percentage activation of SNAP_Switch_ labeled protein in SNAP-tag expressing cells was calculated using the MFI of the samples with or without SNAP-tag pre-treatment and using the following equation:

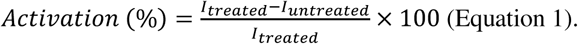

### Fluorescence Microscopy

Imaging by fluorescence microscopy was performed using a 60X 1.3 NA silicone or 40X 0.9 NA air objective with a standard “Pinkel” DAPI/FITC/Cy3/Cy5 Filter set (Semrock). Emission was separated and captured using a 414/497/565/653 nm dichroic mirror and a quad-band bandpass emission filter between 503 – 515 nm and 614 – 804 nm. Images were analyzed with Slidebook 6 (Intelligent Imaging Innovations, Denver, USA). For deconvolution, 10-16 slices were captured with a 0.33 μm step size, exported and deconvolved using the Richard-Lucy algorithm^35,36^ with the CUDA^37^ deconvolution plugin in ImageJ (Version 1.52g, Wayne Rasband, National Institute of Health, USA).

### SNAP_Switch_ Chemical Synthesis

The quenched and attachable benzylguanine substrate was constructed via Fmoc solid-phase peptide synthesis on 2-chlorotrityl chloride resin (200 mg). Fmoc groups were removed using two treatments for 2 minutes and one treatment for 5 minutes with 20% piperidine in DMF. The resin was bubbled for 1 hour in DCM with ethylenediamine (4 equiv, 50 mg, 0.83 mmol). N(α)-Fmoc-N(ε)-azide-L-lysine (1.5 equiv, 0.11 g, 0.3 mmol) was attached with PyBOP (1.5 equiv, 0.16 g, 0.3 mmol) and DIPEA (2 equiv to amino acid, 76 mg, 0.6 mmol) in DMF for 1 hour. A PEG linker was then attached by adding Fmoc-N-amido-dPEG_4_-acid (1.5 equiv, 0.15 g, 0.3 mmol) with PyBOP (1.5 equiv, 0.16 g, 0.3 mmol) and DIPEA (2 equiv, 76 mg, 0.6 mmol) for 30 minutes in DMF. After deprotection, Fmoc-BG-COOH (1.5 equiv, 0.17 g, 0.3 mmol) was attached with PyBOP (1.5 equiv, 0.16 g, 0.3 mmol) and DIPEA (2 equiv, 76 mg, 0.6 mmol) in DMF overnight before washing with DMF, DCM and then followed by deprotection.

15 mg resin was combined in a 1.6 mL Eppendorf tube with the quencher sQSY-21-COOH (0.4 equiv, 5 mg, 5.94 μmol), PyBOP (1.5 equiv) and an excess of DIPEA (10 μL) in anhydrous DMSO for 23 hours, rotating. The resin was transferred back to a solid-phase peptide synthesis column and washed once with DMSO, followed by washes with MeOH until the flow through was clear and then three times with DCM. The resin was dried under N_2_ and the compound cleaved with 4 mL of 50% DCM and 50% HFIP with 10 μL TIPS for 2 hours at room temperature. The mixture was drained into a round bottom flask using HFIP and the solvent evaporated under N_2_. The compound was purified via HPLC with a water/MeCN 0.1% TFA 5 - 100% gradient at a flow rate of 7 mL min^-1^ over 45 minutes. The product mass was confirmed via LCMS.

To the entire fraction from HPLC (4.97 mg), Cy5-NHS (1 equiv) was added with excess DIPEA (10 µL) in 200 μL DMSO and left overnight. The presence of the product mass was confirmed via LCMS and purified using 5 – 100% gradient over 45 minutes at a flow rate of 7 mL min^-1^. The product mass was identified by LCMS, confirmed by HRMS and lyophilized. HRMS (ESI) m/z: calculated for C_108_H_124_N_18_O_25_S_5_ [M + 3H] ^+3^ 745.2602, found 745.2635, calculated for [M + 2H]^+2^ 1117.8882, found 1117.8887, calculated for [M + 2Na]^+2^ 1139.8701, found 1139.8704 (**Supplementary Figure 13**). Purity was estimated by analytical HPLC as ∼ 82% (**Supplementary Figure 14**). SNAP Switch was reconstituted in DMSO to a concentration of 4 mM and stored at −20°C.

### Fluorescence Activation by the SNAP-tag in Solution

The effect of SNAP-tag excess on SNAP_Switch_ was investigated using a fluorescence plate reader. SNAP_Switch_ was initially diluted in PBS to 0.10 μM. 45 μL of this solution was then combined with 0 to 20 molar equivalents of SNAP-tag and PBS to bring the final volume to 65 μL in a 96-well clear bottom black polystyrene microplate. The fluorescence emission at 661 nm was obtained using a 646 nm excitation and was recorded every 10 minutes over 90 minutes total.

### Fluorescence In-Gel Detection

Samples were prepared by combining 5 μM SNAP-tag protein or PBS with 10 µM of SNAP_Switch_ and 1 mM DTT in a final volume of 40 μL. The reaction was incubated at 37°C for 30 minutes then left to cool for 10 minutes before adding 6 μL of 0.2% 2-mercaptoethanol in SDS-page loading dye.^38^ Samples were then heated to 94°C for 2 minutes and added to the wells of a pre-cast 12% polyacrylamide gel (Bio-Rad). The gel was run for 45 minutes at 120 V in running buffer (25 mM Tris, 250 mM glycine, 0.1% SDS at pH 8.3).

### Plasmid Construction

The empty transfection control plasmid pcDNA3.1(-) was purchased from Life Technologies. The TfR-SNAP fusion construct was modified from mEmerald-TFR-20 (a gift from Michael Davidson, Addgene plasmid #54278). Briefly, the sequence encoding SNAP-tag was PCR amplified from pSNAPf (New England BioLabs) with primers to append flanking BamHI and NotI restriction sites (Forward: 5’ – GGA TCC ACC GGT CGC CAC CAT GGA CAA AGA CTG CG – 3’, Reverse 5’ – CGC GGC CGC TTA ACC CAG CCC AGG CTT GCC – 3’). This PCR product was subsequently ligated into mEmerald-TFR-20 at the same sites, replacing the mEmerald coding sequence. A plasmid containing a mTurquoise2-H2A sequence (a gift from Dorus Gadella, Addgene plasmid #36207)^39^ was PCR amplified to append BamHI and NotI restriction sites (Forward: 5’ – ACA GGA TCC ATG GTG AGC AAG GGC GAG GAG – 3’, Reverse 5’ – TGC GGC CGC GTT ATT TGC CTT T – 3’). This PCR product was then ligated into pSNAPf.

Lentiviral transfer plasmids were constructed from pCDH-EF1-MCS-IRES-Puro (System Bioscience). Briefly, the SNAP-tag coding sequence from pSNAPf was subcloned into pCDH-EF1-MCS-IRES-Puro via NheI and BamHI to generate pCDH-EF1-SNAP-IRES-Puro (Cyto-SNAP). The SNAPf-mTurquoise2-H2A, pSNAPf-H2A and TfR-SNAP coding sequences were excised by restriction with NheI and NotI and subcloned into pCDH-EF1-MCS-IRES-Puro. All constructs were verified by DNA sequencing before use.

### SNAP-Tag Expression

The SNAP-tag sequence was PCR amplified from pSNAPf with the following primers, Forward: 5’ – CTGTACTTCCAATCCAATGACAAAGACTGCGAAATGAAGCGCACCAC – 3’, Reverse: 5’ – CCGTTATCCACTTCCAATCCCTCGCAGACAGCGAATTAATTCCAGCA – 3’ and were designed for overlap with pET His6 TEV LIC cloning vector (1B) (a gift from Scott Gradia, Addgene plasmid #29653). The pET His6 TEV LIC cloning vector (1B) was linearized at restriction site SspI and the SNAP-tag sequence was ligated into the plasmid using NEBuilder HiFi DNA Assembly Master Mix. SNAP-tag was subsequently purified by nickel immobilized metal affinity chromatography and used without further modification. Protein concentration was estimated by measuring the UV-vis absorbance at 280 nm and using an extinction coefficient of 22,460 M^-1^ cm^-1^, estimated using ProtParam.^40^

### Estimation of SNAP_Switch_ Concentration

The estimated extinction coefficient of the quenched SNAP-tag substrate was determined using the binary system^41^ Beer-Lambert law: *A*_*n*_ = *ε*_1_*c*_1_*l* + *ε*_2_*c*_2_*l* (Equation 2) where, A_n_ = absorbance at wavelength n, l = path length (cm), εm = extinction coefficient of species m at wavelength n (M^-1^ cm^-1^), c_m_ = concentration of species m at wavelength n (M). Since both the fluorophore and quencher are conjugated to the same molecule, the concentration of each species is the same (c_1_ = c_2_ = c). Using a path length of 0.1 cm, equation 2 reduces to: *A*_*n*_ = 0.1*c*(*ε*_1_ + *ε*_2_) (Equation 3).

The extinction coefficient of each dye at the absorption maximum of the other dye was estimated by measuring the absorbance of a solution with known concentration of Cy5 at 661 nm and of sQSY-21 at 646 nm to give ε_QSY(646)_ and ε_Cy5(661)_. An estimate for the extinction coefficient of SNAP_Switch_ was then obtained using Equation 2 and the absorbance of the conjugate at either of these two wavelengths with the extinction coefficients at the absorption maximum of each dye (ε_Cy5(646)_ = 271,000 M^-1^ cm^-1^ ε_QSY(661)_ = 90,000 M^-1^ cm^-1^).

### Protein Labelling

To label anti-CD44, anti-human transferrin receptor (anti-hTfR) or anti-mouse transferrin receptor (anti-mTfR), 50 μg of protein was diluted in 250 μL of PBS and incubated with 10 equiv. of 1 mg mL^-1^ DIBO-NHS, AF488-NHS or 2 mg mL^-1^ BDP-FL and Cy5-NHS dissolved in DMSO for 2 hours at 4°C. Unreacted DIBO or dye was removed using a 0.5 mL Zeba Spin Desalting Column, 7K molecular weight cut-off, pre-equilibrated with PBS, according to the manufacturer’s instructions. For SNAP_Switch_ labeling,1.25 equiv. of SNAP_Switch_ was added and incubated overnight at 4oC before excess was removed using additional Zeba Spin Desalting Columns.

The degree of labelling (DOL) was estimated by dividing the concentration of SNAP_Switch_ approximated using equation 2 by the protein concentration as determined using the absorbance at 280 nm with the Beer-Lambert law and an extinction coefficient for the protein (ε_antibody_ = 210,000 M^-1^ cm^-1^). The degree of AF488, BDP-FL or Cy5 labelling was calculated using the Beer-Lambert law, the absorbance at 495 nm, 503 nm or 646 nm and an extinction coefficient of ε_AF488_ = 73,000 M^-1^ cm^-1^, ε_BDP-FL_ = 80,000 M^-1^ cm^-1^ or ε_Cy5_ = 271,000 M^-1^ cm^-1^, respectively.

### Oligonucleotide Labelling

The custom 20-mer oligonucleotide modified with dibenzocyclooctyne (DBCO) (Sequence: 5’ - TCA GTT CAG GAC CCT CGG CT – DBCO – 3’) was purchased from IBA Life Sciences, reconstituted in nuclease free water at a concentration of 600 μM and stored at −20°C. The amine modified version (Sequence: 5’ - TCA GTT CAG GAC CCT CGG CT –amino modifier – 3’) was purchased from Integrated DNA technologies and reconstituted using the same conditions. For DBCO-modified oligonucleotide labelling, 1.2 x 10^−8^ mol of the sequence was incubated with SNAP_Switch_ (2 equiv, 24 nmol) and AF488-Azide (1.25 equiv, 15 nmol) overnight at 4°C. Unconjugated dye was spun through 7K molecular weight cut-off Zeba Spin columns until free dye no longer permeated throughout the entire resin.

For amine-modified oligonucleotide labelling, 1.2 x 10^−8^ mol of the sequence was incubated with DIBO-NHS (5 equiv, 60 nmol) for 2 hours at 4°C. Excess DIBO was removed using an Amicon Ultra 0.5 mL Centrifugal Filter with a 3K molecular weight cut-off. Three 5-minute washes were performed using of 500 μL PBS and spinning the units at 13,000 g. SNAP_Switch_ (0.25 equiv) and AF488-Azide (0.25 equiv) were then added and incubated overnight at 4 °C. Unreacted dye was then removed using ethanol precipitation where 1 mL of 9:1 v/v ethanol to 5 M sodium acetate solution was added before spinning the sample for 20 minutes at 4 °C. The supernatant was discarded and 500 μL of −20 °C 70% ethanol was added to the sample without disturbing the pellet. The sample was centrifuged for 20 minutes at 4 °C, the supernatant discarded, and the sample dried by air. The absorbance of the conjugate at 260 nm was measured and degree of labelling was calculated using the previously described extinction coefficients and ε_DNA_= 181,000 M-1 cm^-1^ for the oligonucleotide.

### Anti-Transferrin Receptor Antibody Association

Wild type 3T3 cells or cells stably expressing TfR-SNAP or Cyto-SNAP were seeded in 96-well plates 1 day prior at 30,000 cells per well in 100 μL DMEM supplemented with 10% FBS and penicillin/streptomycin. Proteins were then incubated at with the cells at 5 μg mL^-1^ for 1 to 4 hours before washing twice with PBS and detached with 50 uL TrypLE at room temperature. After 5 minutes, 50 μL 1% bovine serum albumin in PBS was added and the samples were analyzed by flow cytometry.

### Pre-Activated SNAP_Switch_ Conjugates

Wild type 3T3 cells or cells stably expressing TfR-SNAP were seeded in 24-well plates 1 day prior at 100,000 cells per well in 400 μL DMEM supplemented with 10% FBS and penicillin/streptomycin. anti-mTfR dual-labelled with AF488 and SNAP_Switch_ (6.2 μg) was incubated in 1 mM β-mercaptoethanol in PBS (final volume = 93 μL) with or without 30 equiv of SNAP-tag to the SNAP_Switch_ degree of labelling for 1 hour at 37 °C. anti-mTfR was then added to cells at 2.5 μg mL^-1^ and incubated for 1 hour at 37 °C and 5% CO_2_. The cells were washed twice in PBS and detached with 200 μL TrypLE for 5 minutes at room temperature before adding 100 μL 1% bovine serum albumin in PBS. The cells were transferred to a 96-well V-bottom plate and spun at 350 g for minutes. The supernatant was removed, and the cells were resuspended in 100 μL before analysis by flow cytometry.

### Lipofectamine 3000 Induced Endosomal Escape

HEK cells were seeded in 24-well plates 1 day prior at 50,000 cells per well in 400 μL DMEM supplemented with 10% FBS and penicillin/streptomycin. For H2A-mTurquoise-SNAP experiments, cells were transfected with 500 ng DNA or an empty plasmid (pCDNA31(-)) using Lipofectamine 3000, following the manufacturers protocol. 0.75 μL of Lipofectamine 3000 reagent and 1 μL of P300 reagent was used per well. After 16 hours, cells were washed three times in DMEM + 10% FBS before further use. For experiments with TfR-SNAP, Cyto-SNAP and H2A-SNAP the stable cell lines were used.

Cells were transfected with complexes containing either unlabeled oligonucleotide or oligonucleotide labelled with SNAP_Switch_ and Alexa Fluor 488. Samples consisted of 500 ng DNA total with 250 ng each of unlabeled and labeled oligonucleotide (also containing an unlabeled fraction) and were transfected using the same amounts of Lipofectamine 3000 and P3000 previously described. The transfection was performed at 5 time points between 0.5 – 16 hours. For SNAP-tag labeling, the media was replaced with 250 μL DMEM with 10% FBS containing 0.6 μM SNAP-Cell SiR 647 (diluted from 0.6 mM in DMSO). The cells were incubated for 30 minutes, washed three times with DMEM with 10% FBS, then incubated for a further 30 minutes in 400 μL media. All samples were then washed twice in DMEM with 10% FBS and detached using 200 μL TrypLE at 37°C for 5 minutes. After detachment, 100 μL 1% bovine serum albumin in PBS was added to each well and the entire content transferred to a 96-well V-bottom plate. Cells were spun at 350 g for 5 minutes and resuspended in PBS before analysis by flow cytometry.

Cells transfected with H2A-mTurquoise-SNAP and positive for expression were used to compensate for spillover into the AF488 channel. Spillover of mTurquoise and AF488 fluorescence into the Cy5 channel was negligible.

### Antibody Colocalization Imaging

3T3 cells stably expressing TfR-SNAP cells were seeded 1 day prior at 10,000 cells per well in 100 μL, in 96-well plates. Fluorescently labeled anti-hTfR, anti-mTfR or anti-CD44 was then added at 5 μg mL^-1^ and incubated for 1 – 4 hours at 37 °C. For labeling of SNAP-tag at the 4-hour time point, the media was replaced with 100 μL DMEM + 20% FBS containing 5 μM SNAP-Cell 505-Star after 2.5 hours and incubated for 1 hour at 37 oC and 5% CO_2_. The cells were then washed in media three times before incubation for a further 30 minutes. All samples were then washed twice in PBS or three times in FluoroBrite + 10% (for SNAP-tag imagining experiments) and the media replaced with FluoroBrite + 10% FBS with or without Hoechst for 10 minutes prior to imaging.

### Lipofectamine 3000 Endosomal Escape Imaging

HEK293 cells were seeded at 20,000 cells per well in an 8 well chamber slide 1 day prior in 400 μL DMEM supplemented with 10% FBS and 10,000 penicillin/streptomycin. Cells were transfected with 500 ng Cyto-SNAP, H2A-SNAP or an empty plasmid with Lipofectamine 3000 by following the manufacturers protocol. 0.75 μL of Lipofectamine 3000 reagent and 1 μL of P300 reagent was used per well.

After 16 hours, cells were washed 3 times in DMEM + 10% FBS and transfected again with 3100 ng oligonucleotide labelled with SNAP_Switch_ (also containing an unlabeled fraction) and were transfected using the same amounts of Lipofectamine 3000 and P3000 reagent as the previous transfection. The cells were incubated for an additional 16 hours before washing the cells twice with cold FluoroBrite with 10% FBS and then left for 10 minutes on ice. The membrane was then stained with wheat germ agglutinin-AF488 at a final concentration of 1 µg mL^-1^ for 5 minutes before washing an additional two times in FluoroBrite with 10% FBS.

To compare the location of SNAP_Switch_ signal to SNAP-tag labelled with fluorescent SNAP substrates, cells transiently expressing Cyto-SNAP or H2A-mTurquoise-SNAP were treated with 50 µL of 5 µM SNAP-Cell TMR-Star or 3 µM SNAP-Cell SiR 647 in culture media for 30 minutes at 37oC, 5% CO_2_ before washing the cells three times in DMEM with 10% FBS. The media was replaced, and the cells were incubated for a further 30 minutes before replacing the media with FluoroBrite with 10% FBS before imaging.

