## Supporting Information figures are available here. for "SNAP_Switch_: A Molecular Sensor to Quantify the Localization of Proteins, DNA and Nanoparticles in Cells"

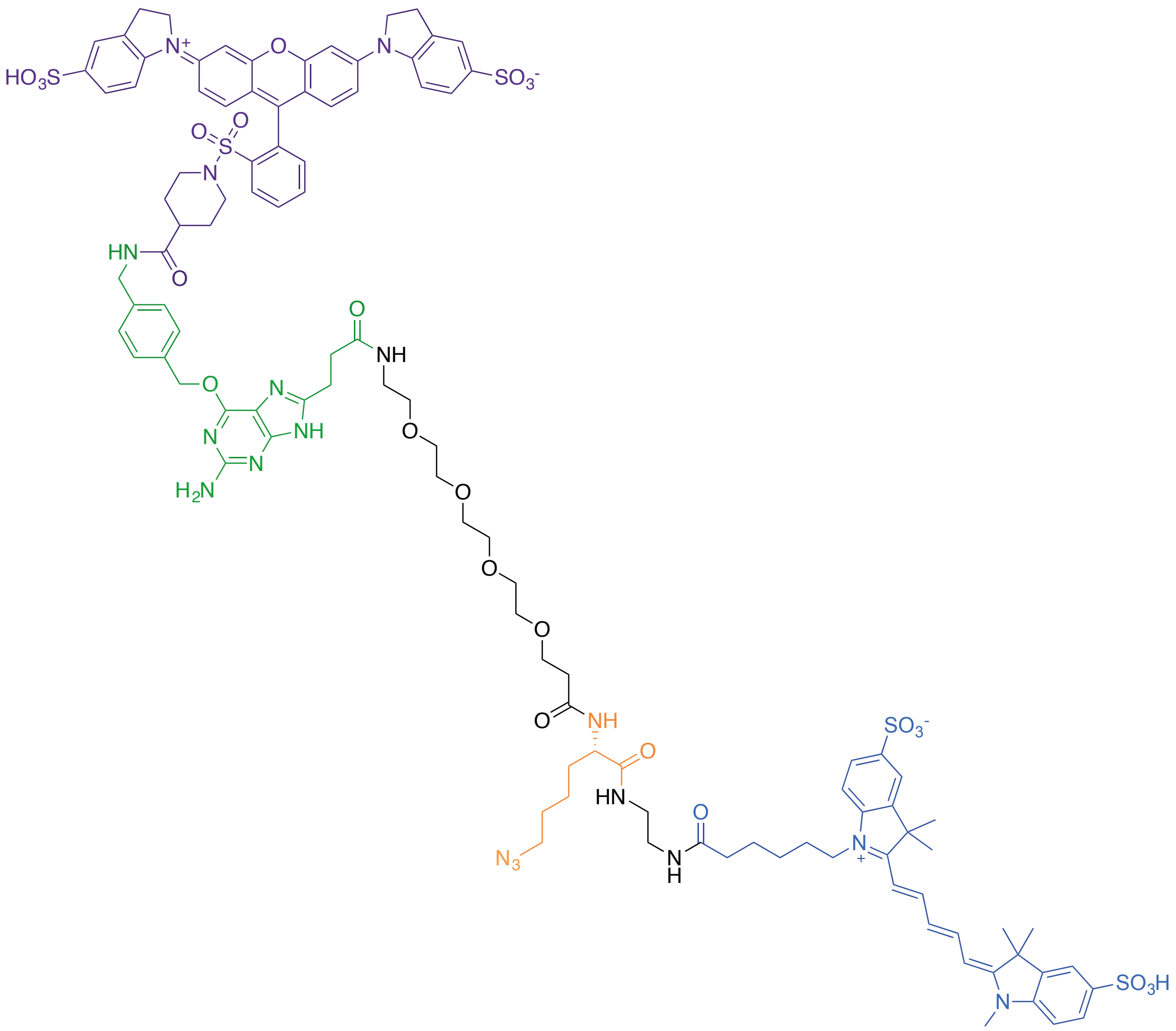

**Supplementary Figure 1** – Chemical structure of SNAP$_{Switch}$**.** QSY-21 (purple) is conjugated to the side of the benzyl guanine substrate (green) that is transferred to the SNAP-tag. The substrate is linked to an azide (orange) for attachment to the particle or protein of interest through a PEG linker. The fluorophore Cy5 (blue) resides on the side of the substrate that remains with the material after interaction with the SNAP-tag.

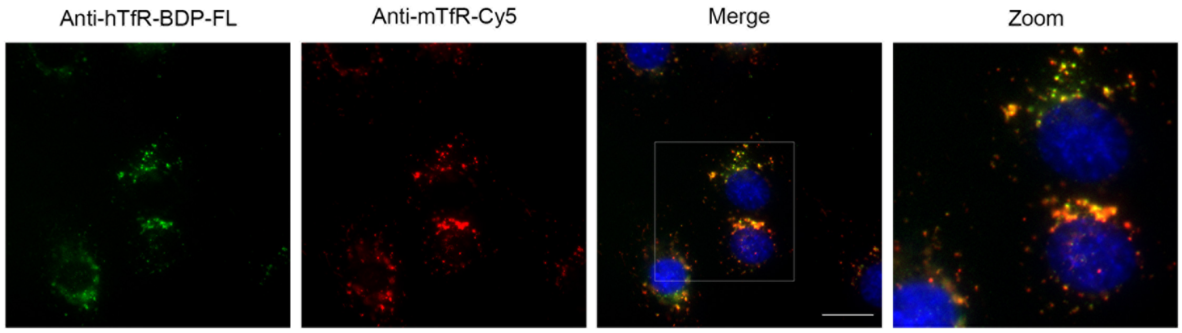


**Supplementary Figure 2** – Deconvolved fluorescence microscopy images of 3T3 cells stably expressing human transferrin receptor fused to SNAP-tag (TfR-SNAP) and incubated with antibodies against human (anti-hTfR) and mouse (anti-mTfR) TfR for 1 hour. Scale bar = 20 µm.

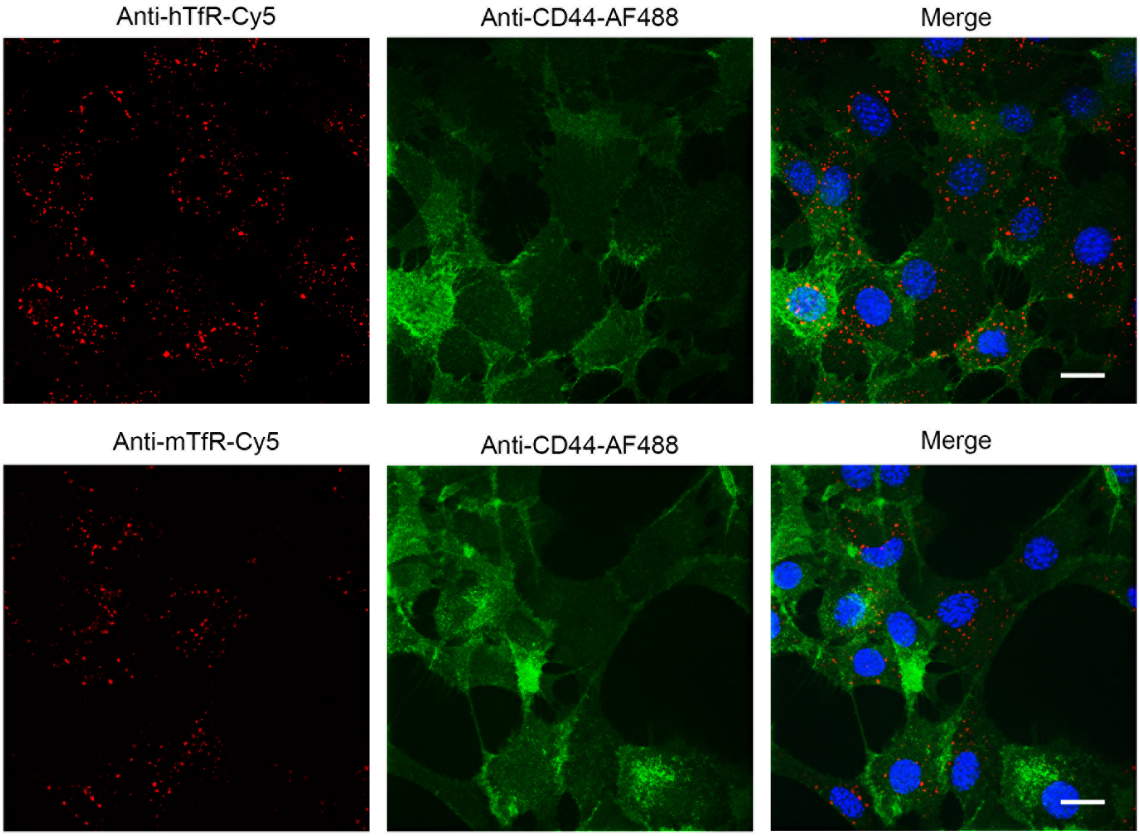


**Supplementary Figure 3** – Deconvolved fluorescence microscopy images of 3T3 cells stably expressing human transferrin receptor fused to SNAP-tag (TfR-SNAP) and incubated with antibodies against CD44, human (anti-hTfR) and mouse (anti-mTfR) TfR for 1 hour. The nucleus is stained with Hoechst. Scale bar = 20 µm.

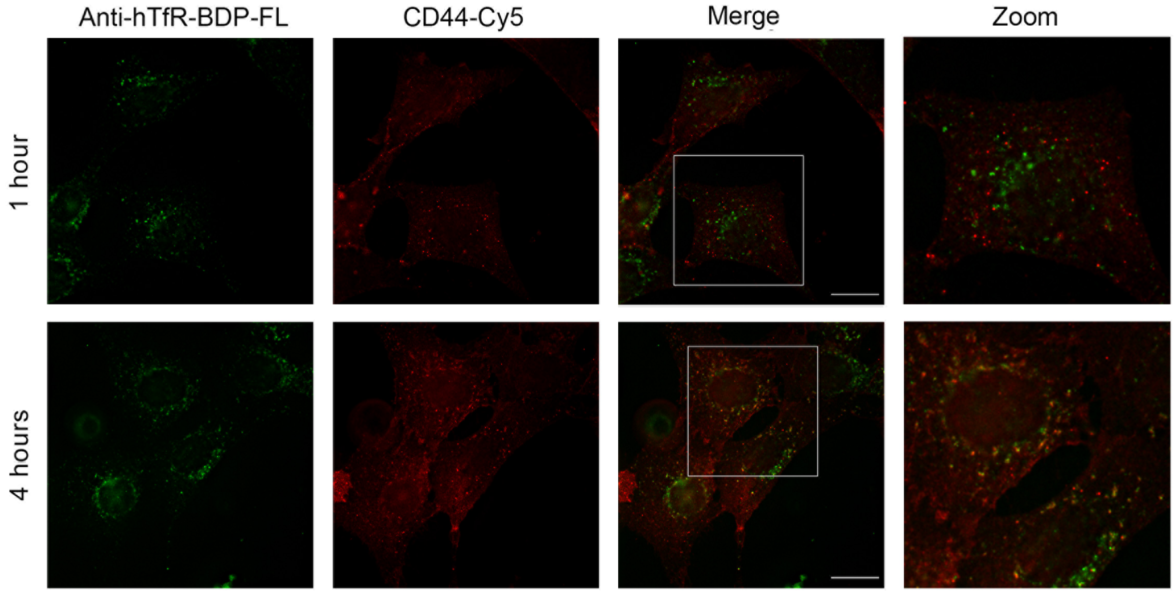


**Supplementary Figure 4** – Deconvolved fluorescence microscopy images of 3T3 cells stably expressing human transferrin receptor fused to SNAP-tag (TfR-SNAP) and incubated with antibodies against CD44 and mouse (anti-mTfR) TfR for 1 or 4 hours. Scale bar = 20 µm.

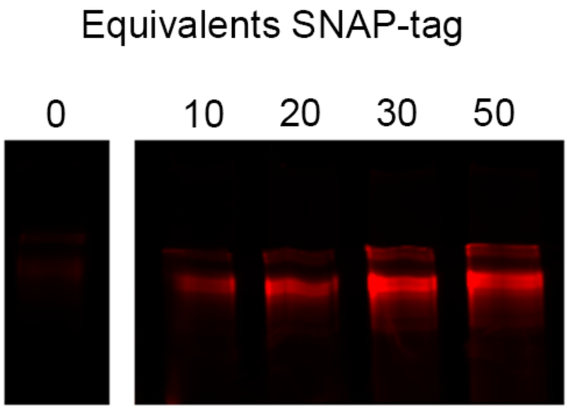


**Supplementary Figure 5** – Fluorescence in-gel detection of anti-mouse TfR antibody (anti-mTfR) labelled with $SNAP_{Switch}$ and incubated with 0 – 50 equivalents of SNAP-tag for 1 hour at 37°C.

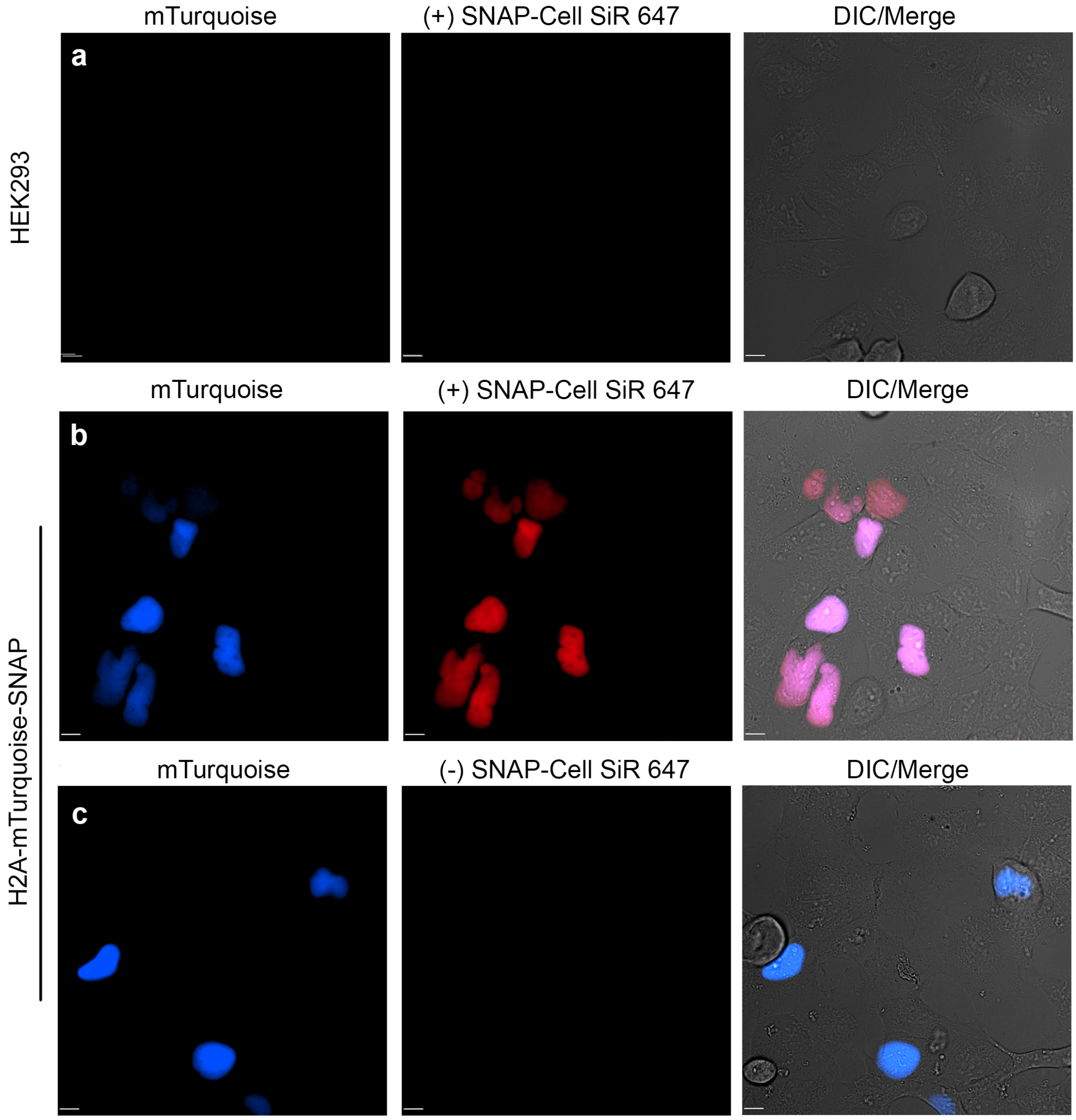


**Supplementary Figure 6** – Fluorescence microscopy images of (**a**) HEK or (**b**) HEK cells transiently expressing H2A-mTurquoise-SNAP, with or (**c**) without treatment SNAP-Cell SiR 647. Brightness and contrast normalized across images, scale bar = 10 µm.

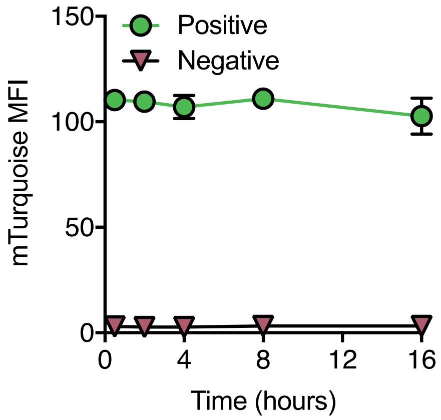


**Supplementary Figure 7** – Flow cytometric analysis of HEK transiently expressing H2A-mTurquoise-SNAP over 16 hours in cells gated for positive or negative expression of mTurquoise. The mean fluorescence intensity is plotted with error bars representing the standard deviation of two experiments in duplicate (n = 4).

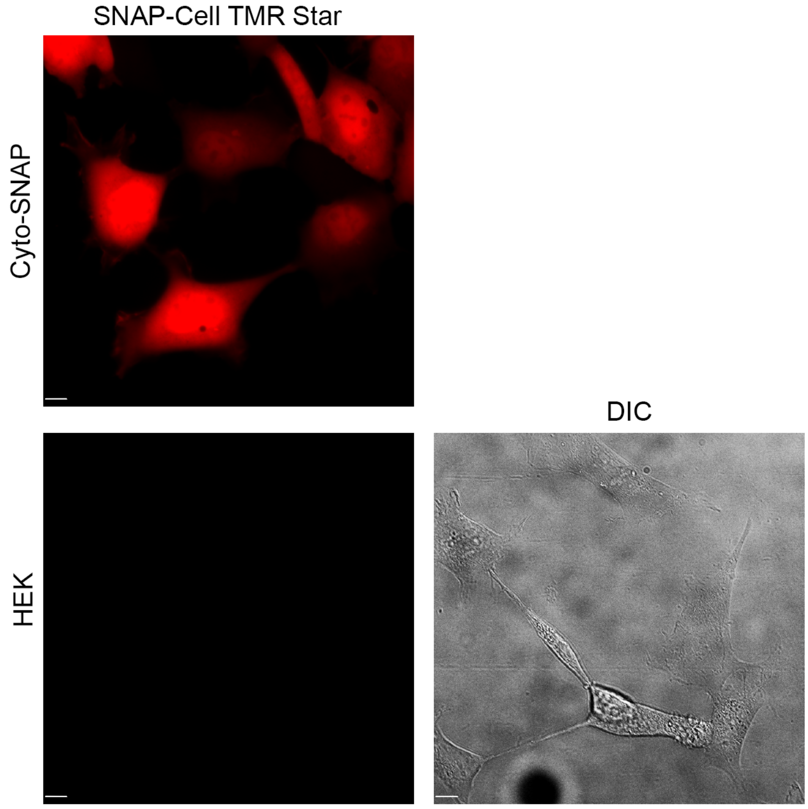


**Supplementary Figure 8** – Fluorescence microscopy images of HEK293 transfected with Cyto-SNAP (top panel) or without transfection (bottom panels) and treated with SNAP-Cell TMR-Star for 30 minutes at 37°C. Brightness and contrast normalized across the fluorescent channel. Scale bar = 10 µm.

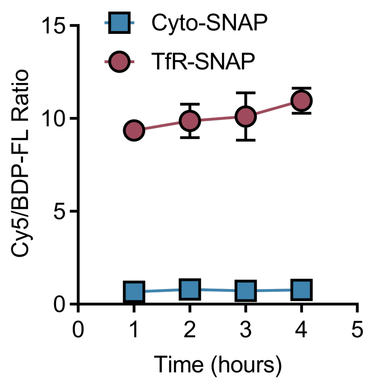


**Supplementary Figure 9 –** SNAP$_{\text{Switch}}$ conjugated to the anti-mouse transferrin antibody anti-mTfR is activated by SNAP-tag fused to the transferrin receptor (hTfR-SNAP) but not enzyme expressed in the cytosol (Cyto-SNAP) over 4 hours. The mean ratio is plotted with error bars representing the standard deviation of two experiments in triplicate (n = 6).

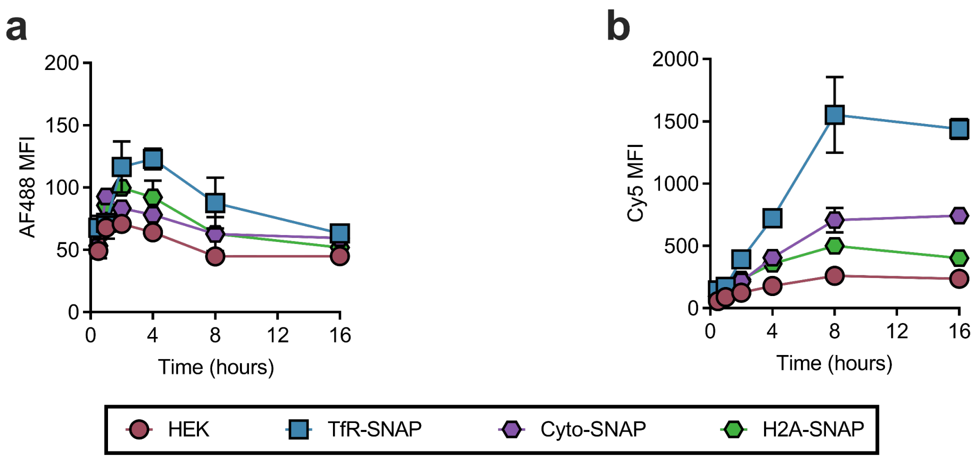


**Supplementary Figure 10 –** HEK cells stably expressing TfR-SNAP, Cyto-SNAP or H2A-SNAP and transfected with Lipofectamine 3000 complexed oligonucleotides labelled with both AF488 and SNAP$_{\text{Switch}}$. (**a**) The association of complexes with cells over time by flow cytometry, measured by the AF488 fluorescence intensity. (**b**) SNAP$_{\text{Switch}}$ signal over time. The mean fluorescence intensity or ratio is plotted with error bars representing the standard deviation, in triplicate (n = 3).

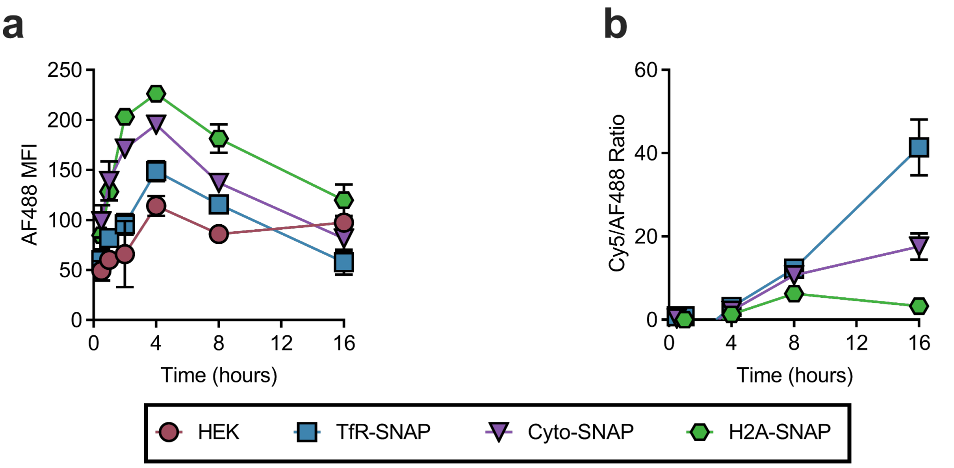


**Supplementary Figure 11** – Repeat experiment of **Figure 3g** in the main text and **SI Figure 12**. HEK cells stably expressing TfR-SNAP, Cyto-SNAP or H2A-SNAP and transfected with Lipofectamine 3000 complexed oligonucleotides labelled with both AF488 and $SNAP_{Switch}$. (**a**) The association of complexes with cells over time by flow cytometry, measured by the AF488 fluorescence intensity. (**b**) The ratio of $SNAP_{Switch}$ to AF488 signal at each time point one experiment in triplicate (n = 3) with the average ratio in HEK cells subtracted from each data point as background.

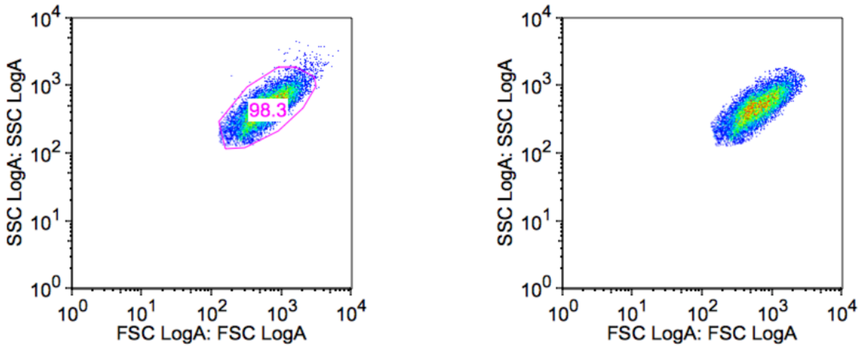


**Supplementary Figure 12** – Example of gating strategy for flow cytometry. Unstained NIH/3T3 cells were gated using the forward versus side scatter log area plot.

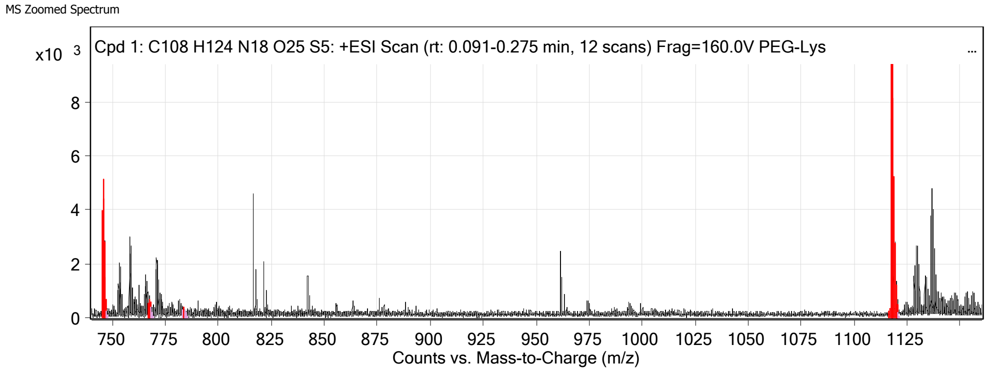


**Supplementary Figure 13** – HRMS $ESI^{+}$ spectra of purified $SNAP_{Switch}$ with found ions highlighted in red.

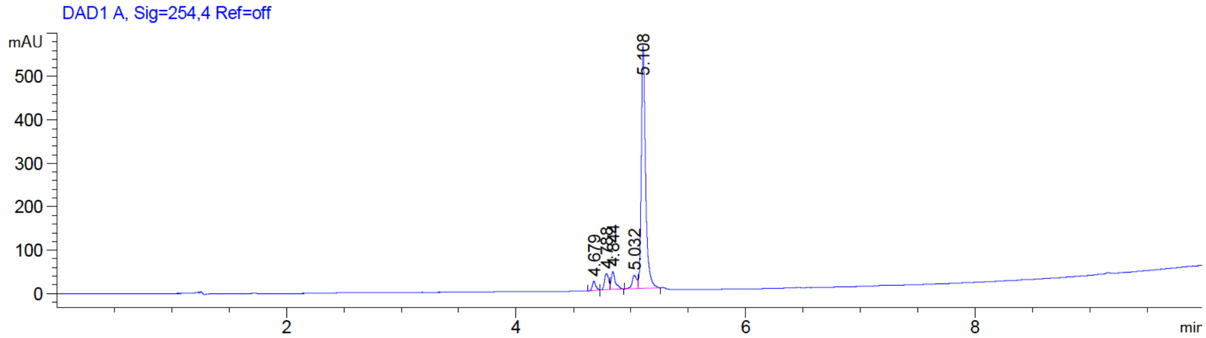


**Supplementary Figure 14** – Analytical HPLC trace of purified SNAP$_{Switch}$. UV absorbance by diode array detector at 254 nm.
